# Multivalent adaptor networks generate nanoscale organisation within T cell signalling condensates

**DOI:** 10.64898/2026.04.28.721251

**Authors:** Shirin Ansari, Pooja Laxman, James Walsh, Zhengmin Yang, Andrea Nunez, Jesse Goyette

## Abstract

Ligand engagement of the T cell receptor (TCR) triggers phosphorylation-dependent assembly of signalling complexes at the plasma membrane, yet how these assemblies are organised to coordinate signal propagation remains unclear. Here, using quantitative super-resolution imaging in primary human T cells, we show that phosphorylated TCR (pTCR) and linker for activation of T cells (pLAT) reside within micrometre-scale signalling condensates that exhibit defined nanoscale organisation. Within these condensates, pTCR and pLAT are spatially interleaved with characteristic spacing of ∼70–100 nm and display short-range dispersion rather than aggregation. Spatial cross-correlation analysis reveals that pLAT is organised around pTCR sites, consistent with a model in which activated TCRs act as local sources of LAT phosphorylation that nucleate condensate assembly. This organisation suggests that multivalent interactions and steric constraints generate structured, yet dynamic, reaction environments that couple receptor activation to downstream signalling. Together, our findings establish that TCR signalling condensates are not amorphous assemblies but possess intrinsic nanoscale architecture and support a general framework in which membrane-proximal signalling is regulated by spatially organised, multivalent assemblies.

## Introduction

Ligand binding to the T cell receptor (TCR) initiates phosphorylation-dependent assembly of signalling complexes at the plasma membrane, mediated by networks of adaptor proteins that couple receptor activation to downstream pathways^1^. While the biochemical interactions underlying these networks are well defined, how they are spatially organised to coordinate signalling reactions remains unclear. Multivalent interactions between phosphorylated receptors, SH2 domain-containing adaptors, and proline-rich region-binding proteins are known to drive the formation of membrane-associated signalling assemblies^2–4^, often described as clusters or condensates^5,6^. However, whether these assemblies represent spatially disordered accumulations of signalling components or possess intrinsic nanoscale organisation has not been resolved. Addressing this question is important for understanding how signalling reactions are coordinated in space within the immunological synapse^7–9^.

The spatial distribution of TCR and LAT has been examined using a range of imaging approaches, leading to differing interpretations of their organisation. Early studies described nanoscale clusters or “protein islands” that reorganise upon activation^10–12^, while others have suggested that TCRs are more uniformly distributed in resting cells^13,14^. More recent work has proposed that signalling occurs within condensate-like assemblies driven by multivalent interactions^6,15^. However, many of these approaches rely on single-molecule localisation microscopy (SMLM) methods that can be affected by fluorophore reblinking and incomplete labelling, complicating quantitative interpretation of spatial organisation^16–18^. These limitations raise the possibility that the nanoscale structure of signalling assemblies may not be fully captured by existing methodologies, and that a quantitative approach capable of resolving both spatial organisation and binding interactions is required.

We previously developed a PAINT-based imaging strategy using recombinant protein domains that bind their targets with defined kinetics (protein-PAINT)^19^. Protein PAINT benefits from the quantitative advantages of PAINT imaging^20^, and overcomes many of the limitations of antibody-based DNA-PAINT strategies, including steric hindrance and underlabelling^18^. In this study, we extend the protein-PAINT framework to revisit the spatial organisation of phosphorylated TCR and LAT in the immunological synapse of primary human T cells activated with cognate peptide–MHC ligands.

In this study, we use quantitative protein-PAINT imaging and analysis to investigate the nanoscale organisation of TCR and LAT signalling assemblies in primary human T cells. By combining probes that report on phosphorylated TCR and LAT with kinetic analysis of binding interactions, we obtain quantitative maps of molecular organisation within signalling condensates. This approach allows us to relate spatial organisation to the underlying multivalent interaction network that drives condensate assembly.

Using these probes, we find that phosphorylated TCR and LAT are distributed within micrometre-scale signalling condensates as spatially dispersed complexes with characteristic nanoscale organisation. Rather than forming tightly aggregated structures, both exhibit regular spacing and short-range dispersion, consistent with a model in which signalling is mediated by spatially distributed, interacting components within condensates. These observations indicate that membrane-proximal signalling assemblies are not defined by molecular clustering alone, but by structured spatial organisation that reflects the underlying multivalent interaction network. Together, our results establish a framework in which TCR signalling occurs within spatially organised condensates that coordinate receptor-proximal signalling through their intrinsic nanoscale architecture.

## Results

### Identification of binding sites

Quantitative protein-PAINT analysis requires accurate identification of individual binding sites from repeated probe binding events. In PAINT imaging, transient probe interactions with a molecular target generate spatially confined clusters of localisations, reflecting repeated binding to the same site. We therefore defined binding sites as regions exhibiting recurrent binding events arising from a single molecular entity or a set of unresolved entities within the localisation precision.

Binding event maps were reconstructed using 5 nm binning, and candidate binding sites were identified as local density maxima and refined by two-dimensional Gaussian fitting to determine centroid positions (Fig S1). Binding events were assigned to a site if they occurred within a 45 nm radius of the centroid, corresponding to approximately three times the localisation precision (Fig S2).

Kinetic parameters were extracted for each site from the temporal sequence of binding events. The apparent association rate (k_on_) was determined from the distribution of off gaps between binding events, and the dissociation rate (k_off_) from the distribution of bound times, as described previously^19,20^.

### Zap70 protein PAINT probe quantitatively images pTCR

In PAINT experiments kinetic rates of probes interacting with regions of the sample carry information of the nature and number of binding sites in this region and can be used for molecular counting experiments^19,20^.

We first measured the bimolecular on rate (k_on_) of the tandem SH2 domains of Zap70 with the membrane distal ITAM of CD3ζ and the ITAM of CD3ε using PAINT imaging of peptides immobilised sparsely on coverslips (Fig 1A-C). Kinetic rates were then fit from the cumulative distribution of off gaps (k_on,_ Fig 1C) and the bound times (k_off_) for binding events from each identified binding site. Measurements of k_on_ (0.86 ± 0.23 µM^-1^s^-1^ for CD3ζ and 0.57 ± 0.22 µM^-1^s^-1^ for CD3ε) made by PAINT were in good agreement with measurements made by SPR (SFig 3, k_on_: 1.1 µM^-1^s^-1^ for CD3ζ) and in our previous work (1.89 ± 0.356 µM^-1^s^-1^ for CD3ζ)^21^.

**Figure 1.**
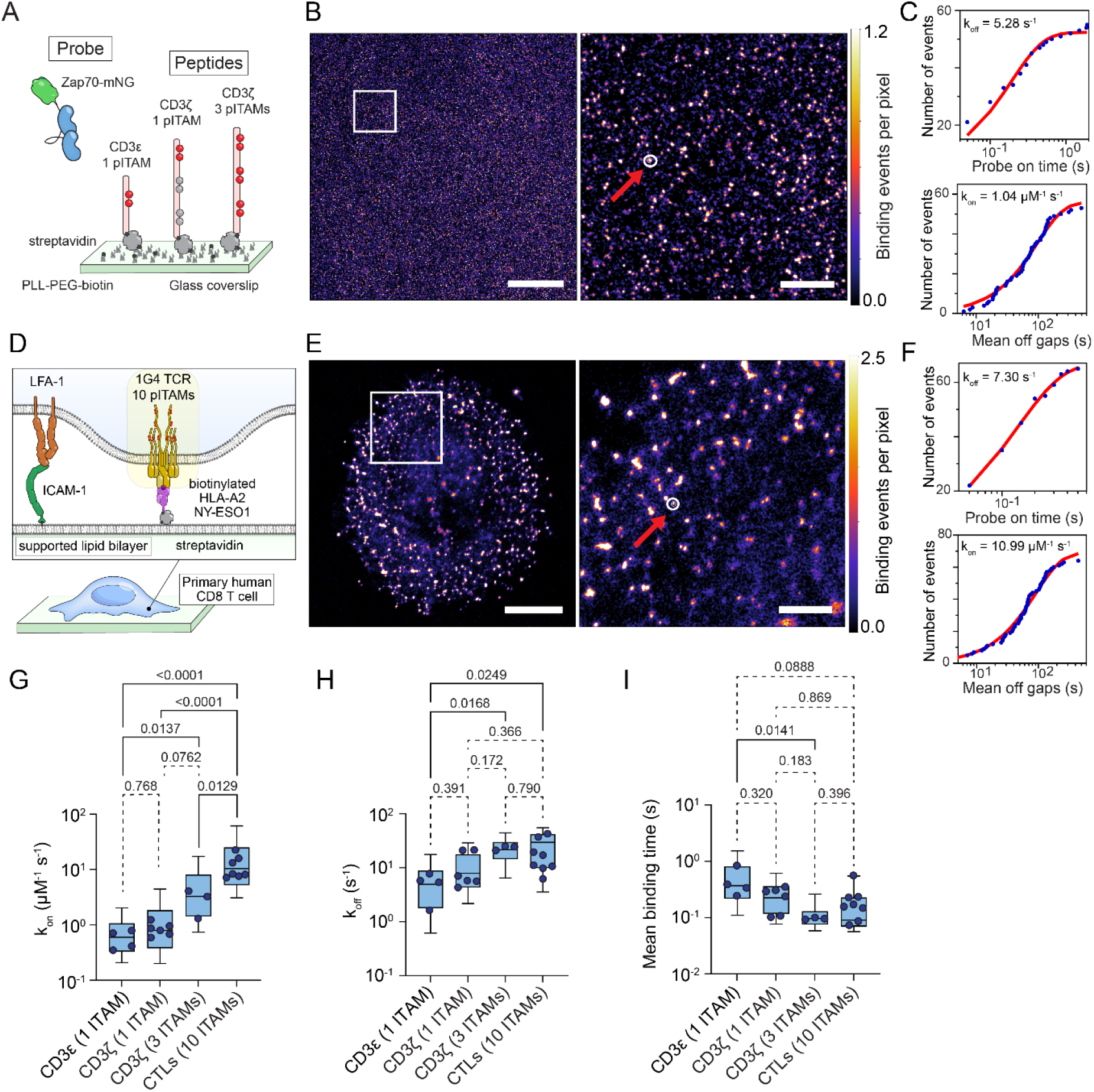
Kinetics of Zap70-tSH2 probe confirm quantitative ITAM quantitation in TCRs *in situ*. (A) Schematic of pPAINT imaging experiments, showing tandem-SH2 Zap70 mNeonGreen (Zap70-mNG) binding to phosphorylated ITAM sites on CD3ζ and CD3ε peptides immobilised on a coverslip. (B) Example reconstructed density image of localised Zap70-mNG binding events on CD3ζ peptide. The red arrow and white circle in the zoom region panel (right) highlights a binding site, defined as a cluster of Zap70 probe rebinding events and identified by 2D gaussian fitting (see methods). Scale bar = 7 μm (left panel) and 1000 nm (right panel). (C) Binding off-rate (k_off_, top panel) and on-rate (k_on_, bottom panel) fit from binding on time and off gap distributions of Zap70-mNG binding to a single peptide binding site identified in (B), right panel (white circle). (D) Schematic of pPAINT imaging experiments for Zap70-mNG binding in primary CD8+ T cells activated on a decorated supported lipid bilayers. (E) Example reconstructed image of localised Zap70-mNG binding events in an activated CD8+ T cell. The red arrow and white circle in the zoom region panel (right) highlights a site of repeated Zap70-mNG rebinding to a phosphorylated TCR. Scale bar = 4 μm (left panel) and 1000 nm (right panel). (F) Single exponential binding off-rate (k_off_, top panel) and on-rate (k_on_, bottom panel) fits from the distribution of on times and off gaps at a single TCR binding site identified in E, right (white circle). (G–I) The distribution of (G) k_on_ (μM^-1^ s^-1^), (H) k_off_ (s^-1^) and (I) mean binding times (s) per binding site. Boxes represent interquartile range, with lines indicating median and whiskers representing 10-90 percentile for all identified binding sites in all images. Individual data points represent global median values per experimental replicate (*in vitro*) or per cell (*in situ*) over three biological replicates. P values from lognormal one-way ANOVA with Tukey’s multiple comparisons tests of the replicate median data points are indicated above each graph.

Where the probe interacts with binding sites that are closer than the localisation precision, the binding on rate (k_on_) of the probe to this region of the imaging field should be linearly proportional to the number of binding sites. In agreement with this principle, when we measured the binding kinetics of our Zap70 probe with a fully phosphorylated CD3ζ cytoplasmic polypeptide the measured k_on_ was 3-fold larger (2.9 ± 1.5 µM^-1^s^-1^) than for single ITAMs peptides (Fig 1G).

Dissociation in single molecule measurements showed complex kinetics, as we previously saw for live cell imaging ^21^ and estimates from single exponential fits are likely a mixture of monovalent and bivalent Zap70-ITAM binding events (Fig S3). The unbinding rate on binding sites with multiple ITAMs in close proximity (fully phosphorylated CD3ζ peptide or TCR in T cells) was also faster than for individual ITAMs (Fig 1H and I), indicating that dynamic Zap70 binding and competition for individual SH2 domains from nearby ITAMs may destabilise the complex in conditions where Zap70 concentration is well below binding saturation for ITAMs^21^.

Having confirmed the kinetics of the probe we next imaged primary human CD8 T cells fixed after 5 min of interacting with supported lipid bilayers presenting ICAM-1 and cognate antigen (Fig 1D). Characteristic clustering of phosphorylated TCRs was observed, as we previously described with this imaging method ^19^. Unlike the peptide samples, where we could control the density and keep peptides sparse enough for minimal overlap, clustering of the TCR in activated T cells led to larger apparent k_on_ rates where TCRs were more closely packed because binding events for binding sites that were within the relatively large 45 nm collection radius were counted multiple times (Fig S4). When we analysed non-overlapping binding sites we found that the k_on_ rate was approximately 10-fold larger (12 ± 6.0 µM^-1^s^-1^) than we measured for a single ITAM, and 3-fold larger than for a fully phosphorylated CD3ζ, consistent with full phosphorylation of all 10 ITAMs in the TCR complex (Fig 1G and Fig S4C).

That the Zap70 protein PAINT probe can detect the full complement of ITAMs on TCRs in the T cell is noteworthy for two reasons. The first is that it contrasts with antibody labelling of affinity tags^18^ and self-labelling tags like SNAP or HALO^22^, which show at best 50% labelling efficiency. The second is that fixation and the presence of endogenous Zap70 does not mask these binding sites for our ZAP70 probe. Taken together this demonstrates that our Zap70 probe provides an unprecedented level of confidence in the localisation and quantitation of phosphorylated TCRs in activated primary human T cells.

### Grb2 protein PAINT probes image pLAT within LAT condensates

We next characterised the binding kinetics of our full length Grb2 probe to phosphopeptides representing the pY171, pY191 and pY226 of LAT, known ligands for the SH2 domain of Grb2^2^, and a proline-rich region (PRR) of SOS1, known to interact with the C-terminal SH3 domain of Grb2^23^ (Fig 2A-C). We also tested the probe in primary human CD8 T cells stimulated by cognate pMHC (Fig 2D-F). Since Grb2 has multiple domains that are important for multivalent interactions that nucleate LAT condensates^6^, we also made probes of the SH2 and C-terminal SH3 domains alone and tested them on peptides and cells (Fig 2A-F).

**Figure 2.**
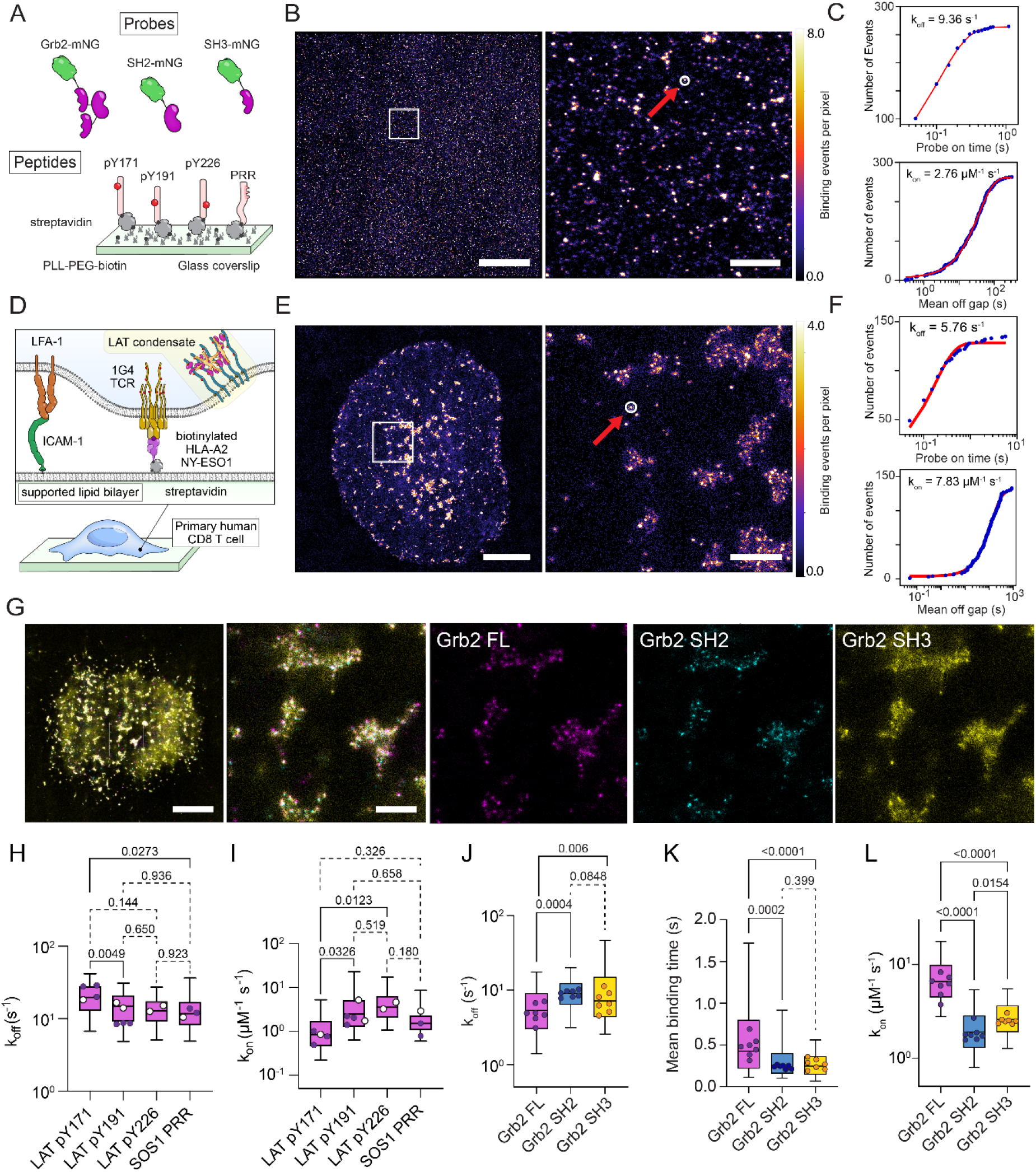
Grb2 pPAINT probes display multimodal binding in primary human T cells. (A) Schematic of pPAINT imaging experiments, showing full length Grb2 mNeonGreen (Grb2-mNG), SH2 domain and C-terminal SH3 domain pPAINT probes. In these experiments synthetic peptides binding partners representing key Grb2 SH2 domain binding sites on LAT (pY171, pY191 and pY226) and a Grb2 SH3 binding site on SOS1 (PRR) were immobilised on glass. (B) Example reconstructed image of localised Grb2-mNG binding events on pY171 peptide. The red arrow and white circle in the zoom region panel (right) highlights a binding site (see methods). Scale bar = 9 μm (left panel) and 1000 nm (right panel). (C) Binding off-rate (k_off_, top panel) and on-rate (k_on_, bottom panel) fit from binding on time and off gap distributions of Grb2-mNG binding to a single peptide binding site identified in (B), right panel (white circle). (D) Schematic of pPAINT imaging experiments for Grb2 pPAINT probes binding in primary CD8+ T cells activated on a decorated supported lipid bilayers. (E) Example reconstructed image of localised Grb2-mNG binding events in an activated CD8+ T cell. The red arrow and white circle in the zoom region panel (right) highlights a site of repeated Grb2-mNG rebinding to a phosphorylated LAT protein. Scale bar = 3 μm (left panel) and 500 nm (right panel). (F) Binding off-rate (k_off_, top panel) and on-rate (k_on_, bottom panel) fit from the distribution of on times and off gaps at a single LAT binding site identified in E, right (white circle). (G) A reconstructed overlay image of multiplexed pPAINT of full length (Grb2-FL, magenta), SH2 domain (Grb2-SH2, cyan) and SH3 domain (Grb2-SH3, yellow). Scale Bar = 3 μm (left panel) and 500 nm (zoom region panels). (H–L) Distribution of (H) k_off_ (s^-1^) and (I) k_on_ (μM^-1^ s^-1^) per binding site from *in vitro* experiments on peptides. Distribution of (J) k_off_ (s^-1^), (K) mean binding times (s) and (L) k_on_ (μM^-1^ s^-1^) per binding site from *in situ* experiments in activated primary CD8+ T cells activated through the TCR. In (H-I) open circles represent measurements from individual domain probes (SH2-mSG for phosphopeptides and SH3-mSG for PRR) and closed circles are full length Grb2-mSG measurements. Boxes represent interquartile range, with lines indicating median and whiskers representing 10-90 percentile for all identified binding sites in all images. Individual data points represent global median values per experimental replicate (*in vitro*) or per cell (*in situ*) over three biological replicates. Open circles in (H) and (I) indicate values from SH2-mNG (pY171, pY191 and pY226) or SH3-mNG (SOS1 PRR), closed circles are values from full length Grb2-mNG. P values from lognormal one-way ANOVA with Tukey’s multiple comparisons tests of the replicate median data points are indicated above each graph.

**Figure 3.**
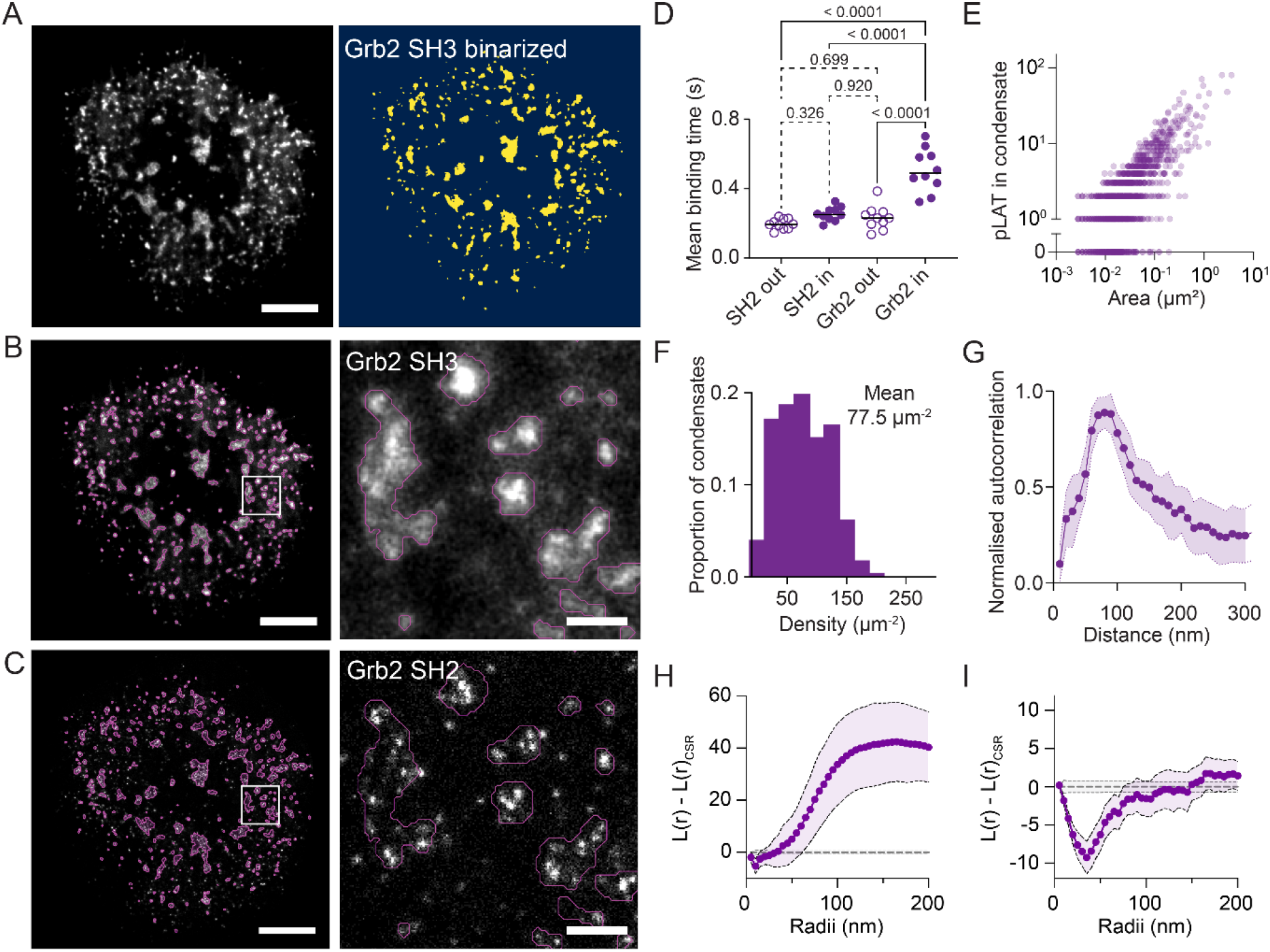
Phosphorylated LAT shows a dispersed and regular organisation within condensates. (A) Reconstructed image of SH3-mNG binding within activated primary CD8+ T cells showing clear micron scale LAT condensates (left panel). An Otsu threshold was used to binarize SH3-mNG images and identify LAT condensate regions (right panel). (B and C) Reconstructed images of (B) SH3-mNG and (C) SH2-mNG binding with LAT condensate regions overlayed as magenta outlines. (D) Mean binding times (s) of full length Grb2 and SH2 probes inside and outside of LAT condensate regions. Each data point represents the mean value for a cell. P values from one-way ANOVA with Tukey’s multiple comparisons tests of the replicate median data points are indicated above each graph. (E) The number of pLAT (see methods for identification), plotted against condensate area. Each data point represents one condensate region. (F) Histogram of pLAT densities within LAT condensates. (G) Circular average spatial auto-correlation of pLAT sites. Data points represent the mean values from 10 cells ± SD (H and I) Ripley’s K functions on entire cell and condensate masked regions. (H) Ripley’s K function with entire cell regions of interest, normalised for variance (L(r)) and for complete spatial randomness (L(r)_CSR_). (I) L(r) – L(r)_CSR_ function restricted to condensate regions. Data points in (G-I) represent means ± 95% confidence intervals of 448 condensates, from 10 cells in 2 biological repeats. Scale bar = 3 μm (left panels) and 400 nm (zoom regions).

Grb2 and the isolated domains had rapid and very similar on and off kinetics on isolated peptides (Fig 2H and I), which agreed well with measurements made using SPR (Fig S5). Only pY171 differed significantly from other peptides, with modestly faster off rate (Fig 2H) and slower on rate (Fig 2I).

When we tested the Grb2 probe in fixed, cognate pMHC-activated, primary human CD8 T cells (Fig 2D), distinct binding sites, visualised as puncta of rebinding events, could be clearly seen (Fig 2E). These binding were clustered at scales of several hundred nanometers in an overall pattern consistent with the size distribution of LAT condensates.

When we performed sequential, multiplex imaging with full length Grb2, isolated SH2 and isolated SH3 domain probes (Fig 2G) we found that the SH2 domain produced clear binding event puncta representing pLAT molecules. The same pLAT molecules were seen in data from the full length probe, but with full length Grb2 there was also diffuse binding events between the pLAT molecules (Fig 2G). In contrast, the isolated SH3 domain probe showed diffuse binding events in regions of clustered pLAT (SH2 binding sites), with regions of higher binding event density that colocalised with individual pLAT molecules (SH2 binding sites) (Fig 2G). The combined pattern of SH2 and SH3 domain binding mirrored that of full length Grb2 probe.

LAT condensates are thought to be held together be weak and dynamic multivalent interactions between LAT, Grb2 and proline rich regions in proteins such as SOS1, SLP76 and ADAP^2,6^. From this understanding of LAT condensates we interpret these images as individual phosphorylated LAT molecules, identified by our Grb2 SH2 probe, interspersed by proteins with long unstructured regions containing large numbers of PRRs, resulting in a cloudlike pattern of low-affinity binding events seen with our Grb2 SH3 domain probe (Fig 2G). Further supporting this interpretation, we found evidence of multivalent binding of our full length Grb2 probe in identified binding sites in cells. Off rates of the full length Grb2 probe were significantly slower (Fig 2J and K), than the individual domains, and the on rate significantly faster, indicating that the Grb2 probe was binding multivalently.

### pLAT is clustered on the micro scale and dispersed on the nano scale

To distinguish pLAT within condensates we created a mask using the density of Grb2 SH3 probe binding to describe regions of high local PRR density (Fig 3A and B). Overlaying the identified condensate regions on data from the Grb2 full length and isolated SH2 probes allowed us to categorise pLAT molecules as falling within or outside of condensates (Fig 3C). As expected, the mean binding time of the SH2 domain probe to pLAT inside and outside was the same, but the full length Grb2 probe had significantly longer binding time on pLAT molecules inside condensates compared with pLAT outside condensates. The large variation in Grb2 probe mean binding times observed in Fig 2K was thus likely due to this mixture of multivalent binding in condensates and SH2 domain binding in pLAT outside condensates.

We next quantified the number of pLAT within the condensates. We observed that the number of pLAT followed a linear trend with respect to condensate area (Fig 3E), with a mean density of ∼80 µm^-2^ (Fig 3F). This relationship was also evident in the raw binding event data, where the influx rate of Grb2 probe binding into condensate regions scaled linearly with condensate area (Fig S6), consistent with a proportional increase in the number of available binding sites.

Phosphorylated LAT within condensates exhibited a non-random spatial distribution, consistent with regular nanoscale spacing. To quantify the organisation, we performed spatial autocorrelation and found a consistent peak between 70-90 nm for pLAT molecules, indicating that there is a regular, repeating pattern of pLAT spacing (Fig 3G). Ripley’s K function has been used extensively to investigate the organisation of molecules within SMLM images, including previous work that suggested the TCR, LAT and other membrane signalling molecules form nanoclusters within the membrane of T cells^24–27^. Ripley’s K function is often normalised to the expected density, such that length scales with clustering are positive and those with dispersion are negative. Ripley’s K analysis is sensitive to large-scale (primary) spatial structure, which can obscure finer-scale secondary organisation unless the analysis is restricted to appropriately defined regions of interest. When we performed Ripley’s K analysis on pLAT with density normalised to the entire cell area, the clustering within condensates dominated the results (Fig 3H). However, if the analysis was restricted to condensate regions, with the expected density normalised for each individual condensate and edge correction applied, a strong dispersion in pLAT at length scales ∼40 nm was observed (Fig 3I).

### pTCR and pLAT are interleaved within LAT condensates

Early SMLM imaging work described TCR and LAT existing in separate “protein islands” that concatenate after activation ^10^. Having found that the organisation of pLAT molecules within condensates was dispersed on the nanoscale we performed multiplex imaging with the Zap70, Grb2 SH2 and Grb2 SH3 probes (Fig 4A) to determine where the pTCR molecules are in relation to LAT condensates. As with the Grb2 multiplex data in Fig 3, we used regions of dense Grb2 SH3 binding to mask regions of LAT condensates (Fig 4A and B). Using this mask, we observed strong association of pTCR within LAT condensates (Fig 4B), with an average density of ∼20 µm^-2^ (Fig 4C). The density of pLAT in LAT condensates in this dataset (∼90 µm^-2^, Fig 4D), which was similar to the density in our previous dataset (Fig 3F), validating the robustness of the analysis and reproducibility between sample preparation. There was also a linear relationship between the number of pTCR and the number of pLAT within LAT condensates (Fig 4E), suggesting that the active Zap70 recruited to pTCRs phosphorylates a consistent amount of pLAT within the condensate.

**Figure 4.**
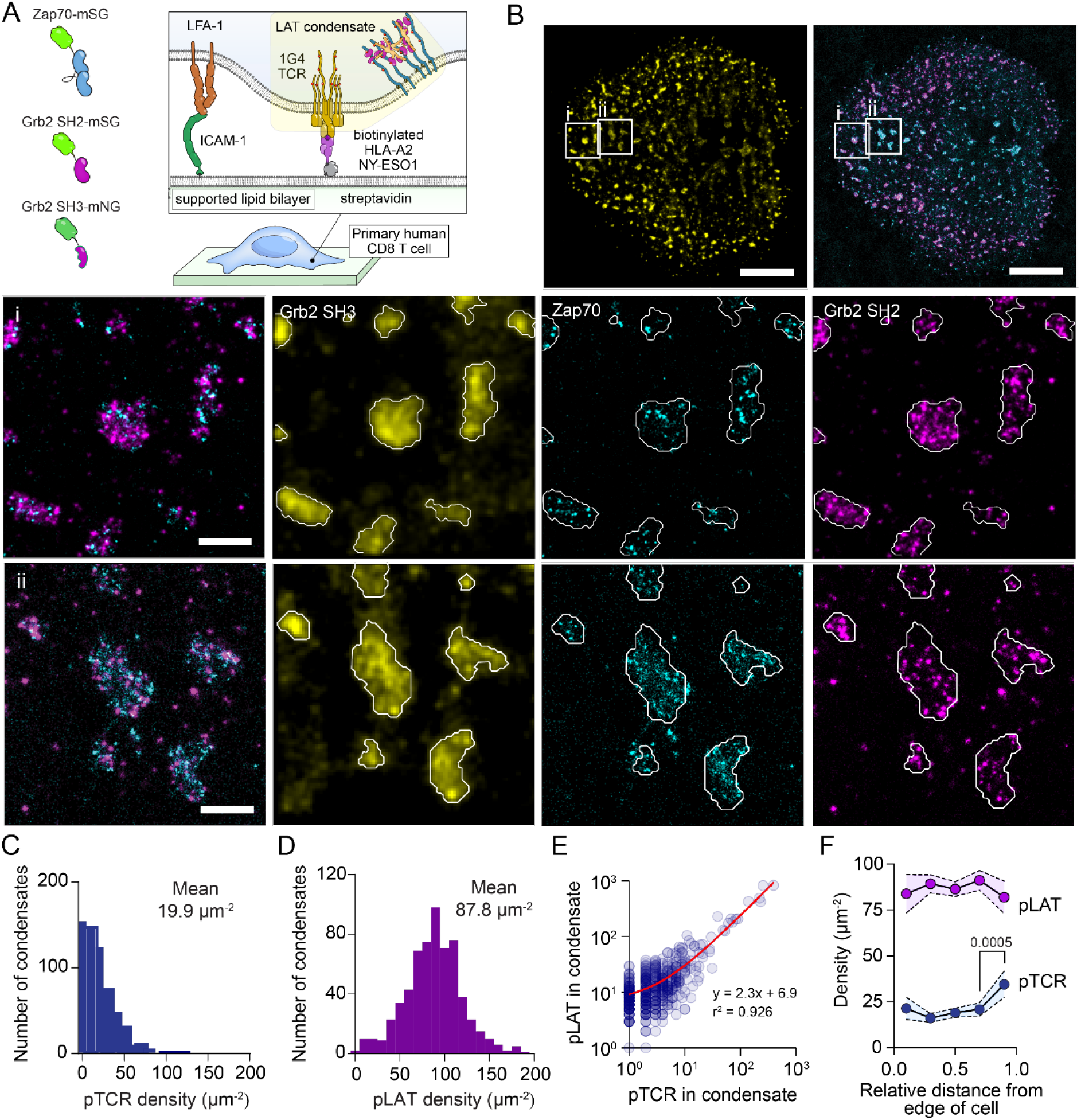
LAT condensates show a consistent pTCR and pLAT density. (A) Schematic of pPAINT imaging experiments for Grb2 pPAINT probes binding in primary CD8+ T cells activated on a decorated supported lipid bilayers. Probes used in these experiments were Zap70 tSH2 mStayGold (Zap70-mSG), Grb2 SH2 domain mStayGold (Grb2 SH2-mSG) and C-terminal SH3 mNeonGreen (Grb2 SH3-mNG). (B) Example reconstructed image of localised Grb2 SH3-mSG (yellow), Zap70-mSG (cyan), Grb2 SH2-mSG (magenta) binding events in an activated CD8+ T cell. Two zoom regions (i) and (ii), showing regions of LAT condensates outlined in white. Scale bar = 4 μm (top panels) and 500 nm (zoom regions). (C and D) Histograms of (C) pTCR and (D) pLAT density within LAT condensates. (D) Number of pTCR plotted against number of pLAT. Each data point represents one condensate. A least squares, linear regression fit (red line) with equation is shown. (F) Density of pTCR and pLAT in condensates plotted against the relative distance from the edge (value of 0) to the centre (value of 1) of the cell. Means ± 95% confidence intervals are shown. P values from one-way ANOVA with Tukey’s multiple comparisons test of values against their neighbouring value are shown (only significant values are shown). Data are from 620 condensates, from 11 cells, in 3 biological repeats.

Activated TCR microclusters are known to form at the periphery of the immunological synapse and migrate to the centre, where they undergo internalisation and dephosphorylation^28^. The density of pLAT and pTCR in condensates remained largely consistent, with a surprising increase in pTCR density observed near the centre of cells (Fig 4F).

Taken together, these analyses indicate that TCR microclusters are not distinct from LAT condensates, but instead represent regions of shared organisation. Reanalysing the data by clustering pTCR using DBSCAN (Fig S7A), as is common for describing TCR microclusters in SMLM imaging^12,29^, revealed the same linear relationship between pTCR and pLAT in pTCR-defined clusters (Fig S7B). This further supports the concept that the terms TCR microclusters and LAT condensates describe the same underlying membrane structure. Across the immunological synapse, the densities of pTCR and pLAT within these regions were largely conserved from the periphery to the centre (Fig S7C). However, pTCR clusters at the cell edge were associated with modestly lower pLAT density and Grb2 SH3 probe influx (Fig S7C and D). This pattern is consistent with a model in which Zap70 recruited to newly engaged TCRs rapidly phosphorylates LAT to nucleate condensates that incorporate pTCR, with these assemblies maintaining a characteristic density as they transit toward the centre of the synapse.

Visual inspection of the reconstructed protein PAINT images suggested that pTCR and pLAT were interleaved with regular spacing (Fig 5A), which was particularly evident when examining small regions of the reconstruction (Fig 5B). Autocorrelation analysis revealed characteristic peak distances of approximately 70 nm for pTCR (Fig 5C) and 90 nm for pLAT (Fig 5D), indicating that both species exhibit nanoscale organization within condensates. In contrast, spatial cross-correlation analysis between pTCR and pLAT displayed a monotonically decaying correlation (Fig 5E). Nearest neighbour analysis supported the interpretation that a small length scales (<50 nm) pLAT and pTCR are more likely to be found near each other than to another homotypic species (Fig 5F).

**Figure 5.**
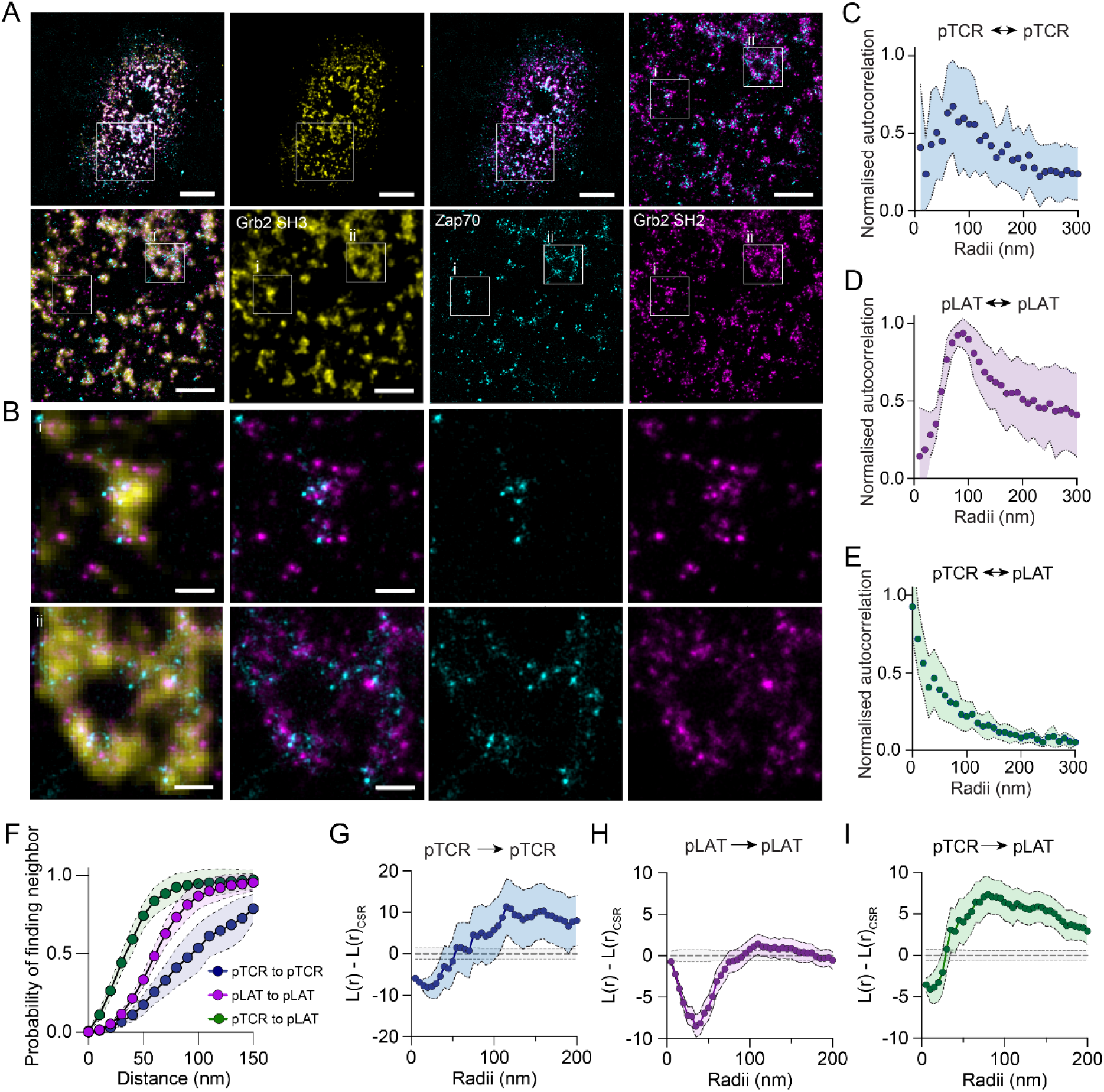
Organisation of phosphorylated LAT and phosphorylated TCR within LAT condensates. (A) Example overlay image of localised Grb2 SH3-mNG (yellow), Zap70-mSG (cyan), Grb2 SH2-mSG (magenta) binding events in an activated CD8+ T cell. Two zoom regions (i) and (ii), showing regions of LAT condensates outlined in white. Scale bar = 3 μm (top panels) and 1000 nm (zoom regions, right and bottom panels). (B) Enhanced zoom regions (i) and (ii) as indicated in zoom regions of (A). Scale bar = 200 nm. (C–E) Circular average of two dimensional spatial auto-correlation of (C) pTCR, and (D) pLAT and (E) cross-correlation of pTCR and pLAT. Data points represent the mean values from 11 cells ± SD. (F) Nearest neighbour analysis of pTCR and pLAT within condensate regions. Cumulative probability distribution of finding at neighbour plotted against increasing search radius for indicated pairs. (G–I) Variance and complete spatial randomness-normalized Ripley’s K function (L(r) – L(r)_CSR_) restricted to condensate regions for (G) pTCR to pTCR, (H) pLAT to pLAT, and (I) pLAT to pTCR. Data points in (F) represent means ± 95% confidence intervals of 11 cells, in 3 biological repeats and (G–I) represent means ± 95% confidence intervals of 620 condensates, from 11 cells, in 3 biological repeats.

Ripley’s K analysis further showed short-range (∼25 nm) dispersion for pTCR (Fig 5G) and dispersion on a slightly larger scale (∼40 nm) for pLAT (Fig 5H). In contrast, cross-Ripley’s analysis between pTCR and pLAT revealed dispersion only at very small separations (peak dispersion at 10-15 nm), consistent with molecular exclusion effects, but clustering at ∼100 nm (Fig 5I), suggesting that pLAT is organized around pTCR sites in a structured manner rather than forming a spatially random mixture.

Together, these results indicate that the condensates possess internal organization, possibly maintained by competing forces that both promote coalescence and preserve nanoscale spacing between molecular species. This implies that while pTCR acts as a nucleating site for pLAT phosphorylation and condensate assembly, the resulting structures are not amorphous but spatially patterned by intermolecular interactions and steric constraints.

## Discussion

The spatial organisation of membrane-proximal signalling has been widely described in terms of clusters or condensates^5,6,10,11^, yet how these assemblies are internally structured to coordinate signalling has remained unclear. Here, we define the nanoscale architecture of T cell signalling condensates and show that phosphorylated TCR and LAT are not randomly distributed or tightly aggregated, but instead exhibit regular spacing and spatial interleaving within micrometre-scale assemblies. This organisation is consistent with a model in which activated TCRs act as local sources of LAT phosphorylation that nucleate condensate formation^15,30^, generating structured environments in which signalling components are positioned with defined nanoscale relationships. These findings indicate that multivalent adaptor networks do not simply drive condensation, but establish spatially organised reaction environments in which the balance of multivalent interactions and steric constraints governs molecular arrangement^3,4,6^. In this view, signalling condensates are not amorphous assemblies, but structured and dynamic platforms that couple receptor activation to downstream signalling through their intrinsic spatial organisation.

The distributions of pTCR and pLAT within condensates show characteristic spacings of ∼70–90 nm and consistent short-range dispersion, distinct from either random mixing or tight aggregation. Such regularity likely arises from the interplay of multivalent attraction and steric repulsion among signalling components. Activated TCRs act as local sources of LAT phosphorylation^15,30^, which nucleate the assembly of condensates through multivalent interactions between pLAT, Grb2, and PRR-containing adaptors such as SOS1, SLP76, and ADAP^2–4^. A plausible explanation for the observed nanoscale structure is that the extended, intrinsically disordered domains of these scaffold proteins impose an entropic repulsion^31,32^ counterbalancing the polyvalent SH2-phosphotyrosine and SH3-PRR binding that keeps the condensate together, preventing molecular collapse and maintaining defined spacing. In this model, the resulting condensates are cohesive but porous, sustaining dynamic accessibility for kinases and phosphatases while preserving spatial order.

These findings reconcile two frameworks that have often been considered distinct: the TCR microcluster^33^, widely described in imaging studies^28,34^; and LAT condensates, also termed LAT clusters or signalosomes but redefined as dynamic assemblies following the work of Su et al.^6^. Rather than separate entities, they represent different views of the same underlying assembly observed from different perspectives. The nanoscale interleaving of pTCR and pLAT may be important for signalling fidelity, allowing efficient kinase–substrate coupling^35^ while preventing unrestrained lateral propagation.

Protein-PAINT affords a kinetic and stoichiometric precision that overcomes major artefacts of single-molecule localization microscopy. By directly quantifying binding-site density and kinetics, it avoids overcounting due to imperfect accounting for fluorophore reblinking^16^ and undercounting from incomplete labelling^18^. The ability to detect all phosphorylated ITAMs on individual TCRs, and to map multivalent binding interactions of Grb2 within condensates, reveals nanoscale order that was previously invisible. These methodological advances support a reappraisal of what constitutes “clustering” in super-resolution datasets, emphasising the need to distinguish between true molecular proximity and statistical artefacts.

Beyond TCR signalling, similar structured condensates may underlie other membrane-proximal pathways that rely on multivalent scaffolds, including B cell receptor and receptor tyrosine kinase systems^36,37^ (Case et al., 2019; Lin et al., 2023). The principle that weak, reversible interactions generate mesoscale organisation without loss of molecular accessibility may represent a general design rule for signalling hubs based on multivalent pY-SH2 and PRR-SH3 interaction networks.

Our findings refine the spatial model of early TCR signalling by demonstrating that LAT condensates possess intrinsic nanoscale organisation rather than forming amorphous assemblies. By resolving the interleaved arrangement of phosphorylated TCR and LAT within these structures *in situ* in primary human CD8 T cells, this work bridges mesoscale observations from imaging studies ^28,34^ with biochemical and *in vitro* reconstitution ^6^ approaches to multivalent signalling. Importantly, the ability to quantify binding site density and organisation at molecular resolution reveals structural features that are not accessible by conventional imaging or bulk assays. These results position signalling condensates not simply as phase-separated compartments, but as spatially structured reaction environments shaped by the interplay of multivalent interactions and physical constraints. Understanding how this nanoscale organisation evolves over time and contributes to signal propagation, sensitivity, and decision-making will be an important direction for future studies of T cell activation.

## Methods

### Peptides and Plasmids

**Table 1.** List of plasmids constructs used to generate cell lines.

| Plasmids | Details |
| --- | --- |
| pHR-1G4 | NY-ESO-1 specific TCR (1G4 TCR) alpha and beta subunits in lentiviral pHR-SIN-BX vector. |

**Table 2.** List of Peptides used for *in vitro* protein-PAINT. CD3ϵ and LAT pTyr peptides were obtained from Thermofischer Scientific and SOS1 PRR-Cy5 was obtained from GeneScript.

| Peptides | Sequence |
| --- | --- |
| CD3ε ITAM (Biotinylated) | biotin-Ahx-GQNKERPPPVPNPD [pY] EPIRKGQRDL [pY] SGLNQRR |
| CD3 $\zeta$ single phosphorylated ITAM | biotin-Ahx-GKGHDLL [pY] QGLSTATKDT [pY] DALHMQ |
| CD3 $\zeta$ three phosphorylated ITAMs | Cytoplasmic region of full length CD3 $\zeta$ with N-terminal hexa-histidine and Avitag expressed in *E.coli*, biotinylated with recombinant BirA labelling kit and phosphorylated with a Streptag II-Y505F mutant Lck fusion expressed in HEK 293T cells and purified with Strep-Tactin beads (IBA Life Sciences). |
| LAT pY171 (Biotinylated) | biotin-Ahx-SAFSMESIDD [pY] VNVPESGE |
| LAT pY191 (Biotinylated) | biotin-Ahx-AEASLDGSRE [pY] VNVSQELH |
| SOS1 Proline Rich Region-Cy5 | Cy5-GGPAPSIDRSTKPP |

#### Peptide Synthesis

Peptide sequences were fused with an N-terminal hexahistadine tag and an Avitag and cloned into a pTRC vector backbone. For the singly phosphorylated ITAM construct, the tyrosines in the first two ITAMs were mutated to alanines. Peptides were recombinantly produced in *Escherichia coli* BL21 cells. Biotin (100 μM) was added to media at the time of induction with IPTG (500 μM) to drive the biotinylation of the Avitag by endogenous BirA. Following cell lysis by sonication and clarification by centrifugation (10 000 xg for 15 min), the CD3ζ peptides was purified by Ni^2+^-affinity chromatography and subsequent phosphorylated *in vitro* using recombinant Lck.

#### Cell culture

CD8+ T cells were isolated from peripheral mononuclear blood cells (PBMCs). Human buffy coats were obtained from Australian Red Cross lifeblood (ethics: HC210396). Samples were diluted 1:2 in RPMI 1640 (Gibco) before centrifugation separation of peripheral mononuclear blood cells (PMBC) by Ficoll-paque Plus procedure. After PMBC collection, cells were washed (PBS, 2 mM EDTA, 0.5 % (w/v) BSA) and then purified using human CD8+ T Cell Isolation Kit (Miltenyi Biotec) following manufacturer’s instructions. Resulting cells were cryopreserved in foetal bovine serum with 10 % (v/v) dimethyl sulfoxide.

CD8+ T cells were maintained in RPMI medium supplemented with 10% (v/v) fetal bovine serum (FBS), 2mM L-glutamine, 1mM penicillin and streptomycin, 25mM HEPES, 50μM β-Mercapthoethanol, 2mM Sodium Pyruvate (all from Invitrogen) and 25ng/mL of IL-2 (Australian Biosearch). Cells were cultured in suspension at 37^°^C and 5% CO_2_ at a cell density of 1×10^6^ to 2×10^6^ cells per mL.

HEK293T cells were cultured in DMEM (Gibco) media containing phenol red and 2mM glutamine, this was supplemented with 10% FBS, 1mM penicillin and streptomycin (all from Invitrogen). Cells were cultured adherently at 37^°^C and 5% CO_2_ at a confluency of 80%. After which cells were trypsinised with 0.05% Trypsin (Sigma-Aldrich) for 5 minutes at 37°C, washed in DMEM and passaged to 30% confluency in fresh DMEM.

#### Lentiviral transduction to generate 1G4 TCR expressing CD8+ T cells

HEK293T cells were cultured for at least 1 week after revival from cryovials prior to lentiviral transduction. On the day before transfection, HEK293T cells were passaged to a confluency of 30-40% in a T-75 flask to ensure the cells would reach a 70-80% confluency. On the day of the transfection, a plasmid cocktail containing 7.5μg of pRSV, 7.5μg of pGAG, 2.9μg of pVSV-G, 6.3μg pHR-1G4 TCR and 48μg Polyethylenimine (Sigma-Aldrich), was made in serum-free DMEM. Media was replaced with serum-free DMEM and to the cells, the plasmid mix was added. This was incubated at 37°C for 5 hours. After which, the serum-free media was discarded and was replaced with complete DMEM media and incubated at 37^°^C. Supernatant containing virus particles were collected at 24, 48 and 72 hours and filtered through a 0.45 μm filter.

On the day of transduction, after the virus was collected at the 72-hour mark, the total viral supernatant was centrifuged for 90 minutes (16,000 rpm, 4°C) to pellet the virus. The supernatant was then gently decanted, and the pellet resuspended in 1mL of complete RPMI media. The concentrated viral supernatant was added to CD8+ primary T cells (prepared at 2×10^6^ cells in 1mL) in a 12-well plate. To the cell culture, 25 μL of CD3/CD28 Dynabeads (Gibco) were added for every 1 million cells to activate them and 25 ng/mL IL-2. The cells were incubated at 37°C, 5% CO2 for 48 hours, after which cells were washed with fresh media to remove the virus. The cells were incubated for another 3 days before the removal of the beads using the Dynabead separation magnetic rack (ThermoFisher Scientific). Cells were once again washed with fresh complete RPMI media and incubated for 24 hours prior to use.

#### Preparation of coverslips and imaging chambers

Glass coverslips were cleaned by sequential sonication for 10 minutes each in ethanol, ultrapure water, and 1M NaOH, followed by thorough rinsing in ultrapure water before being dried. The coverslip was then treated in an air plasma cleaner (Harrick Plasma) for 5 minutes. After plasma treatment, either an 8-well removable chamber (Ibidi) for peptide imaging, or a PDMS device for multiplex imaging in cell using microfluidic flow cells was attached to the coverslip. PDMS device fabrication was adapted from ^38^, with flow channels measuring 8 mm in length, 2 mm in width and 0.15 mm in height.

For imaging on the Active Stabilisation Feedback microscope ^39^, 3μm polystyrene beads (Spherotech) diluted 1:200 in ultrapure water were added to each well. The beads were then dried onto the coverslip by incubation at 70°C for 30 minutes. Once dried, the chamber was plasma cleaned again and prepared for either support lipid bilayer formation for cell activation or for peptide immobilization for *in vitro* PAINT. For microfluidic chamber experiments, all subsequent solutions were pulled through the device at a flow rate of 50 μL min^−1^.

#### Generation of Liposome and preparation of supported lipid bilayer (SLB)

A lipid mixture with a final composition of 96.5% DOPC, 2% DOGS-NTA, 1% biotin-DOPE, and 0.5% DOPE-PEG5000 (mol%; Avanti Polar Lipids) was prepared. The lipids were combined to dried overnight under vacuum to create a lipid film. The film was rehydrated in ultrapure water to a final lipid concentration of 1 mg/mL for at least 30 minutes, then extruded 15–17 times through a 0.1 μm membrane filter to produce homogeneous liposomes. Extruded liposomes were stored at 4°C under nitrogen gas for up to one week.

To generate SLBs, liposomes were diluted to 0.2 mg/mL in ultrapure water, and 10 mM CaCl_2_ was added. The suspension was added to plasma-treated chambers and incubated for 30 minutes at room temperature to allow bilayer formation on the glass surface. The resulting bilayer was gently washed with ultrapure water and incubated with 10 mM EDTA for 10 minutes, followed by the addition of 10 μg/mL streptavidin and 1 mM NiCl_2_ for 15 minutes. Finally, the wells were washed with 1% BSA/PBS and ligands added to decorate the bilayers.

#### Cell activation

To activate 1G4 TCR-expressing CD8+ T cells, 1 µg/mL of biotinylated pMHC (HLA-A2-SLLMWITQV) (MBL Life Science), and 500 ng/mL of His-tagged ICAM-1 (ACROBiosystems) were added to wells in 1% BSA/PBS for 1 hour. The wells were thoroughly washed with 1% BSA/PBS to remove unbound ligands. Pre-warmed media was added and the sample placed into the incubator to equilibrate prior to addition of cells. Approximately 200,000 - 250,000 cells in 50 μL were added per well and incubated for 4 – 10 minutes at 37°C to allow activation. For fixation, pre-warmed 4% paraformaldehyde (PFA) was added and incubated for 15 minutes at 37°C. Samples were then washed with 1x PBS and blocked overnight in 5% BSA/PBS at 4^°^C.

#### Peptide preparation for imaging

Plasma-treated chambers were coated with 0.1 mg/mL PLL-PEG-Biotin for 20 minutes and then washed with 1x PBS. Streptavidin (10 µg/mL) was added for 20 minutes and subsequently washed with 1x PBS after. Biotinylated peptides, diluted to 10nM in 1x PB, were then added to each well for 20 minutes to allow binding to streptavidin. Excess peptides were washed with 1x PBS.

#### Protein-PAINT probe preparation and total internal reflection microscopy (TIRF) imaging

Protein-PAINT probes were recombinantly produced in *E. coli* as previously detailed previously^19^. Probes were prepared in imaging buffer containing 1x PBS, 1% BSA, 0.1% saponin, 1 mM DTT, and 1 mM EDTA. Probes were diluted in imaging buffer to final concentrations ranging from 1 nM (for cell imaging) to 10 nM (for *in vitro* peptide imaging). For multiplexed imaging, wells were washed with imaging buffer between probes until previous probe signal was completely removed.

Images were acquired on the custom-built 3D-stabilised active-feedback microscope ^39^. The microscope was configured with a 100x oil-immersion TIRF objective (Nikon; numerical aperture 1.49) and two cameras for simultaneous fluorescent and brightfield acquisition. Fluorescent signals were captured on an Orca-Fusion C14440 CMOS camera (Hamamatsu) with a 2048 × 2048-pixel sensor and an effective pixel size of 86 nm. Laser power was adjusted according to experimental requirements, typically ranging from 5-80 mW for the 488 nm channel and 50mW for the 647 nm channel (Vortran). Imaging acquisition length ranged from 25,000 to 400,000 frames with an exposure time of 50ms.

For active stabilisation, the 3-μm polystyrene beads were used to lock the field of view and provide real-time drift correction during image acquisition. The beads were illuminated using an 850 nm infrared LED and imaged using a Manta GigE infrared camera (Allied Vision) at 20 ms exposure, producing a 1936 × 1216-pixel field of view with a 58 nm pixel size.

#### Multiplexed imaging

Sequential protein-PAINT imaging was performed by exchanging probes within the same imaging chamber. After acquisition with each probe, chambers were washed extensively with imaging buffer until residual fluorescence signal returned to baseline levels. Complete removal of prior probe signal was confirmed by the absence of detectable binding events over at least 1,000 frames.

Probes were applied sequentially at concentrations of 1–10 nM in imaging buffer. The order of probe addition was varied between experiments to control for potential artefacts arising from probe binding or sample perturbation.

#### Image processing

Raw fluorescent image stacks were analysed using the Picasso software package ^40^ to obtain information regarding the localisations of each single binding event. Each fluorescent spot in the image frames was analysed by fitting it a 2D gaussian distribution to generate a sub-pixel localised x,y coordinate. Localised .hdf5 files were analysed using Picasso’s Render program. Here localisations were linked into individual binding events by linking detections appearing within a radius of 80 nm over consecutive imaging frames, with a 5-frame gap joining length and drift corrected using redundant cross-correlation. These localisations were saved as .hdf5 files for further analysis. To analyse the saved localisations, we have developed a novel data analysis pipeline to allow for the kinetic studies of these localisations.

#### Protein-PAINT kinetic analysis and binding site identification

Single-molecule localisation data obtained from protein PAINT image series using Picasso, and linked into binding events, as detailed above, were analysed using a custom Python-based package: Python Image Processing Environment (PIPE), available at GitHub (DOI to be included in final publication).

Binding sites were identified from reconstructed binding event density maps (5 nm pixel size) by automated detection of local maxima followed by two-dimensional Gaussian fitting. Candidate sites were accepted based on amplitude and width thresholds consistent with the experimentally determined localisation precision (typically 10–15 nm). Individual binding events were assigned to binding sites if they fell within a 45 nm radius of the fitted site centre, corresponding to roughly three times the localisation precision.

Kinetic parameters were extracted from the temporal sequence of binding events associated with each site. The dissociation rate (k_off_) was obtained from the distribution of bound times (dwell times) using maximum likelihood estimation assuming exponential or exponential decay models where appropriate. Association rates (k_on_) were determined from the distribution of dark times (off gaps) between consecutive binding events at each site, normalised by probe concentration. For sites with overlapping assignment regions, apparent increases in k_on_ were interpreted in the context of event sharing between neighbouring sites.

All fitting procedures were performed using custom scripts in PIPE, using scipy and numpy libraries.

#### Condensate segmentation and masking

LAT condensate regions were defined using the spatial distribution of Grb2 SH3 domain probe binding events. Reconstructed images (10 nm pixel size) were smoothed using a Gaussian filter (σ = 1 pixel) and thresholded using Otsu’s method to generate binary masks. Thresholding was applied on a per-cell basis to account for variability in overall signal intensity.

Binary masks were further processed to remove small objects below 0.01 μm^2^ and to fill holes within condensates. The resulting masks were used to classify binding sites and localisations as either inside or outside condensate regions.

Condensate area was calculated directly from the binary mask, and all spatial analyses restricted to condensates were performed within these masked regions.

All masking, binding site assignment to masked regions and calculations were performed using custom scripts in PIPE.

#### Spatial statistics analysis

#### Autocorrelation and cross-correlation analysis

Spatial autocorrelation of binding site distributions was computed using a fast Fourier transform (FFT)-based approach implemented in the PIPE analysis software. Binding site maps were converted into reconstructed images with 10 nm binning, and the two-dimensional autocorrelation function was calculated via Fourier convolution. Radial averaging was performed to obtain one-dimensional autocorrelation profiles as a function of distance.

Autocorrelation functions were normalised to values at zero spatial shift to normalise for density differences between datasets. Spatial cross-correlation between pTCR and pLAT binding sites was computed in the same way as autocorrelation, but with separate images generated for each species.

#### Ripley’s K function

Ripley’s K function was calculated for binding site coordinates using custom Python scripts implemented in the PIPE software package. The variance-stabilised L(r) function was computed as L(r) = sqrt(K(r)/π), and results are presented as L(r) – L(r)_CSR_, where CSR denotes complete spatial randomness.

Edge effects were corrected using an isotropic correction method. For analyses restricted to condensates, K functions were calculated within individual condensate masks and normalised by the local density within each condensate.

Cross-Ripley’s K analysis was performed to quantify spatial relationships between species. The cross-K function was calculated using pairwise distances between pTCR and pLAT sites and normalised to the expected distribution for independent spatial processes with matched densities.

#### Nearest neighbour analysis

Nearest neighbour distances were computed for all binding site pairs using Euclidean distance metrics. Cumulative distribution functions were generated for homotypic (pTCR–pTCR, pLAT–pLAT) and heterotypic (pTCR–pLAT) nearest neighbour distances.

#### DBSCAN clustering analysis

Clustering of pTCR binding sites was performed using the DBSCAN algorithm as implemented in the scikit-learn Python package. Binding site centroid coordinates were used as input. Parameters were set to a search radius (ε) of 200 nm and a minimum of 3 points per cluster.

Clusters identified by DBSCAN were used to define pTCR-enriched regions for comparison with LAT condensates. Cluster properties, including area and number of binding sites, were calculated from the spatial extent of clustered points.

#### Density and stoichiometry analysis

Binding sites identified from protein-PAINT data were treated as individual molecular entities. Molecular density was calculated as the number of binding sites per unit area within defined regions (either entire cell footprint or condensate masks).

For condensate analyses, density was calculated per condensate and reported as mean values across all condensates. Relationships between molecular counts (e.g., pTCR vs pLAT) were assessed using linear regression.

#### Surface plasmon resonance (SPR)

Binding kinetics of protein-PAINT probes were independently measured by surface plasmon resonance using a Biacore S200 instrument. Biotinylated peptides were immobilised on streptavidin-coated CM5 sensor chips, and probe proteins were injected at multiple concentrations in running buffer (10 mM HEPES pH 7.4, 150 mM NaCl, 3 mM EDTA, 0.005% Tween 20). Association (k_on_), dissociation (k_off_) rates and equilibrium binding constants (K_D_) were obtained by fitting sensorgrams to a 1:1 Langmuir binding model using GraphPad Prism.

#### Precision and resolution estimation

Localisation precision was estimated using the Nearest Neighbour (NeNa) method^41^. Image resolution was determined by Fourier ring correlation (FRC) analysis of reconstructed localisation datasets^42^. Both analyses were performed using established implementations within the Picasso and PIPE software frameworks.

#### Statistical analysis

All statistical analyses were performed using GraphPad Prism. Data distributions were assessed for log-normality where appropriate. Comparisons between groups were performed using one-way ANOVA with Tukey’s multiple comparisons test unless otherwise stated.

Data are presented as median with interquartile range unless otherwise indicated. Individual data points represent biological replicates (cells or experiments as specified in figure legends).

## Code availability

Code used to analyse data and prepare images in this work is available from https://github.com/goyette-lab/PIPE

## Acknowledgements

We thank the generous funding support from the National Health and Medical Research Council (NHMRC) Ideas Grants (GNT2021307 and GNT1183588 to J.G.). We would like to acknowledge the support and help from the staff of Katharina Gaus Light Microscopy Facility (KGLMF) and the Flow Cytometry Facility at the Mark Wainwright Analytical Centre, UNSW, specifically Dr Emma Johansson Beves and Dr Elvis Pandzic.

