## Supplemental Figures for "Multivalent adaptor networks generate nanoscale organisation within T cell signalling condensates"

### Supplementary material

#### Supplementary Figure 1

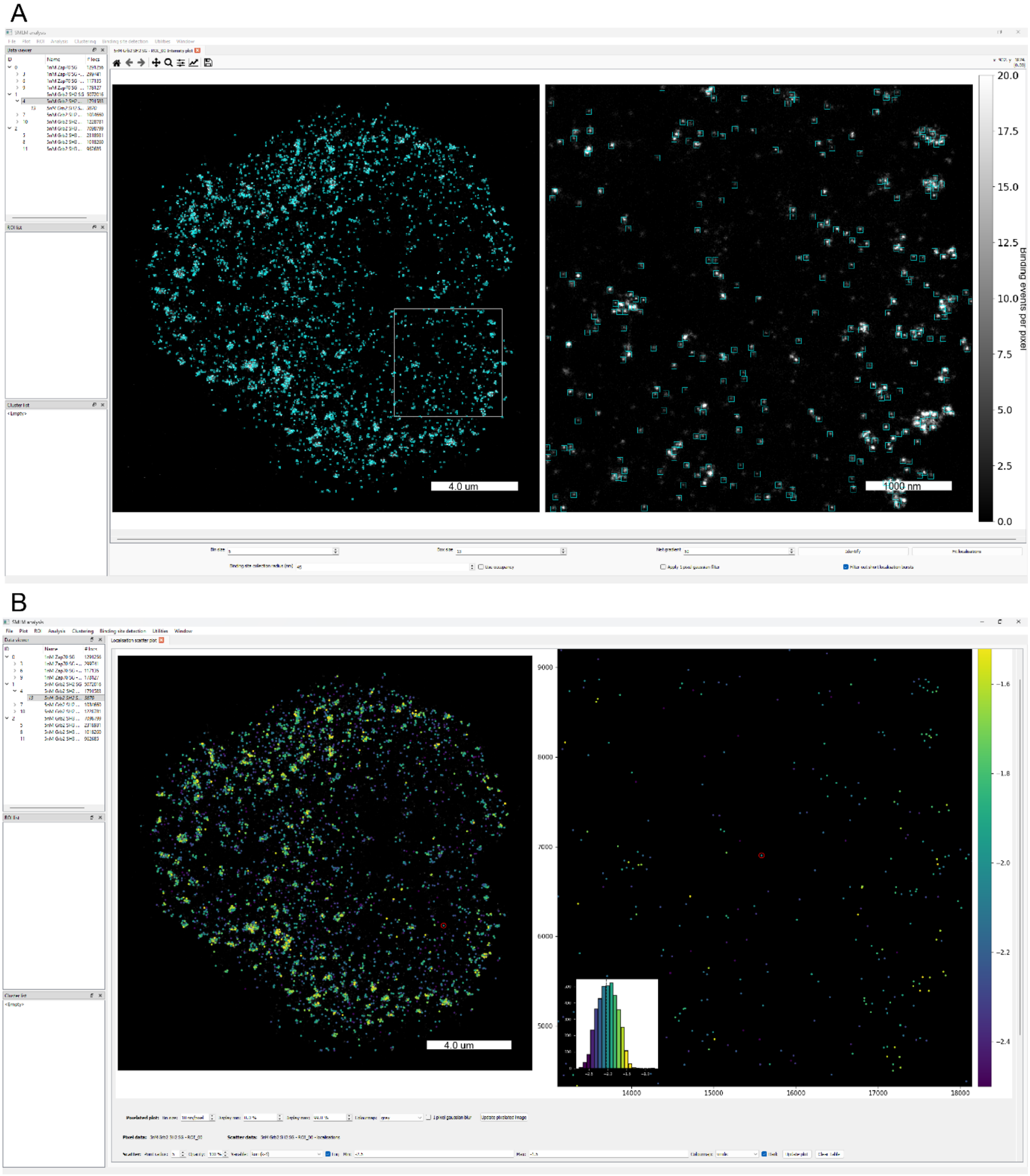


**Figure S1: Automated identification of binding sites**

(A) Binding sites were identified in reconstructed PAINT images using a 2D guassian fitting algorithm. Displayed is an example screenshot from Python Image Processing Environment (PIPE) software showing identification of binding sites (cyan boxes) for ZAP70 protein PAINT probe.

(B) Identified ZAP70 probe binding sites pseudo-coloured by binding on-rate.

**Supplementary Figure 2**

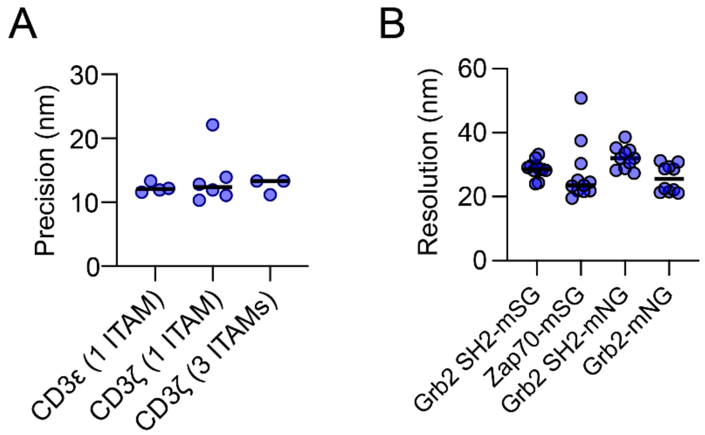


**Figure S2: Estimate of precision and resolution in protein PAINT data.**

(A) Precision estimated by the NeNa method from Zap70-mNG/peptide datasets shown in main text Fig 1.

(B) Resolution of reconstructed PAINT images from linked localization data binned at 5 nm per pixel using the Fourier ring correlation method.

#### Supplementary Figure 3

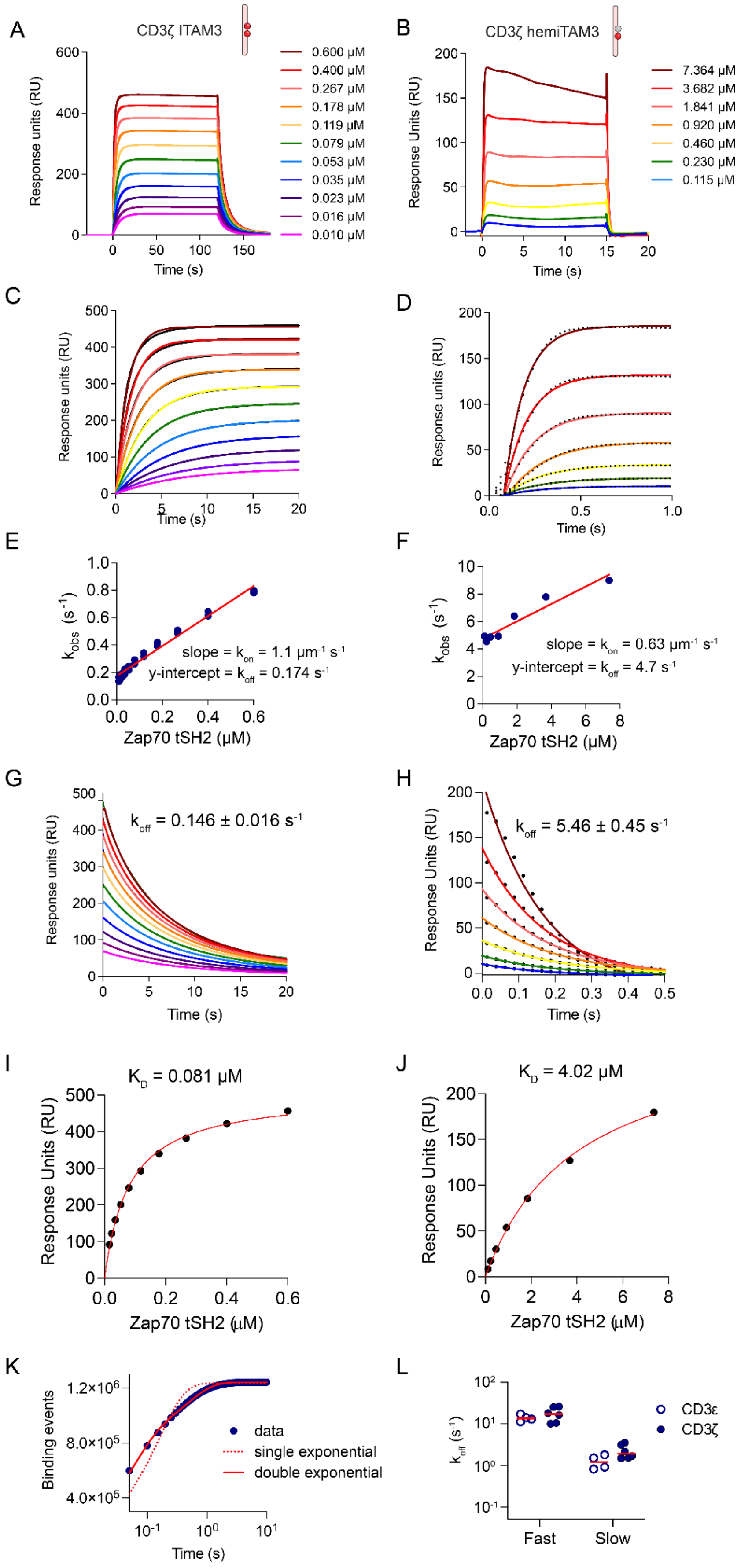

**Figure S3: Binding kinetics of ZAP70 PAINT probe**

(A and B) Raw overlayed SPR traces of ZAP70-mNG probe on a biphosphorylated ITAM peptide (A) and a hemi-phosphorylated ITAM peptide (B) with a sequence identical to the membrane distal ITAM of CD3ζ. Concentrations of probe Zap70 probe injected are shown.

(C and D) Association rate ($k_{obs}$) fits of a simple 1:1 binding model from the association phase of Zap70 probe on a biphosphorylated ITAM peptide (C) and a hemi-phosphorylated ITAM peptide (D).

(E and F) Association rates ($k_{obs}$) vs ligand concentration with linear regression fits for a biphosphorylated ITAM peptide (C) and a hemi-phosphorylated ITAM peptide (D). As expected from previous work[1], the $k_{on}$ rate for hemiphosphorylated ITAM peptide was half that of the biphosphorylated ITAM peptide, and $k_{off}$ was much lower for biphosphorylated taget demonstrating bivalent interaction.

(G and H) Dissociation constants ($k_{off}$) fit from the dissociation phase of SPR traces for biphosphorylated ITAM peptide (G) and a hemi-phosphorylated ITAM peptide (H) agreed with results the $k_{obs}$ fits (E and F).

(I and J) Equilibrium binding and $K_D$ fits from data in (A and B).

(K) Compared with the ensemble measurements of SPR, single molecule measurements reveal more complex binding modes. Shown are cumulative sums of binding event lifetimes in an example experiment with Zap70 probe binding to a biphosphorylated ITAM. A double exponential model (solid red line) fits the data much better than a single exponential model (dotted line) indicating at least two binding modes are evident in the single molecule binding data.

(L) Fast and slow component rates for the double exponential fits of cumulative sums of binding event lifetimes in experiments on phosphorylated CD3ε ITAM peptide and the membrane distal ITAM (ITAM3) of CD3ζ are shown. Rapid transient single SH2-phosphotyrosine interactions together with longer-lived, photobleaching-limited, bivalent interactions are evident in single molecule data.

**Supplementary Figure 4**

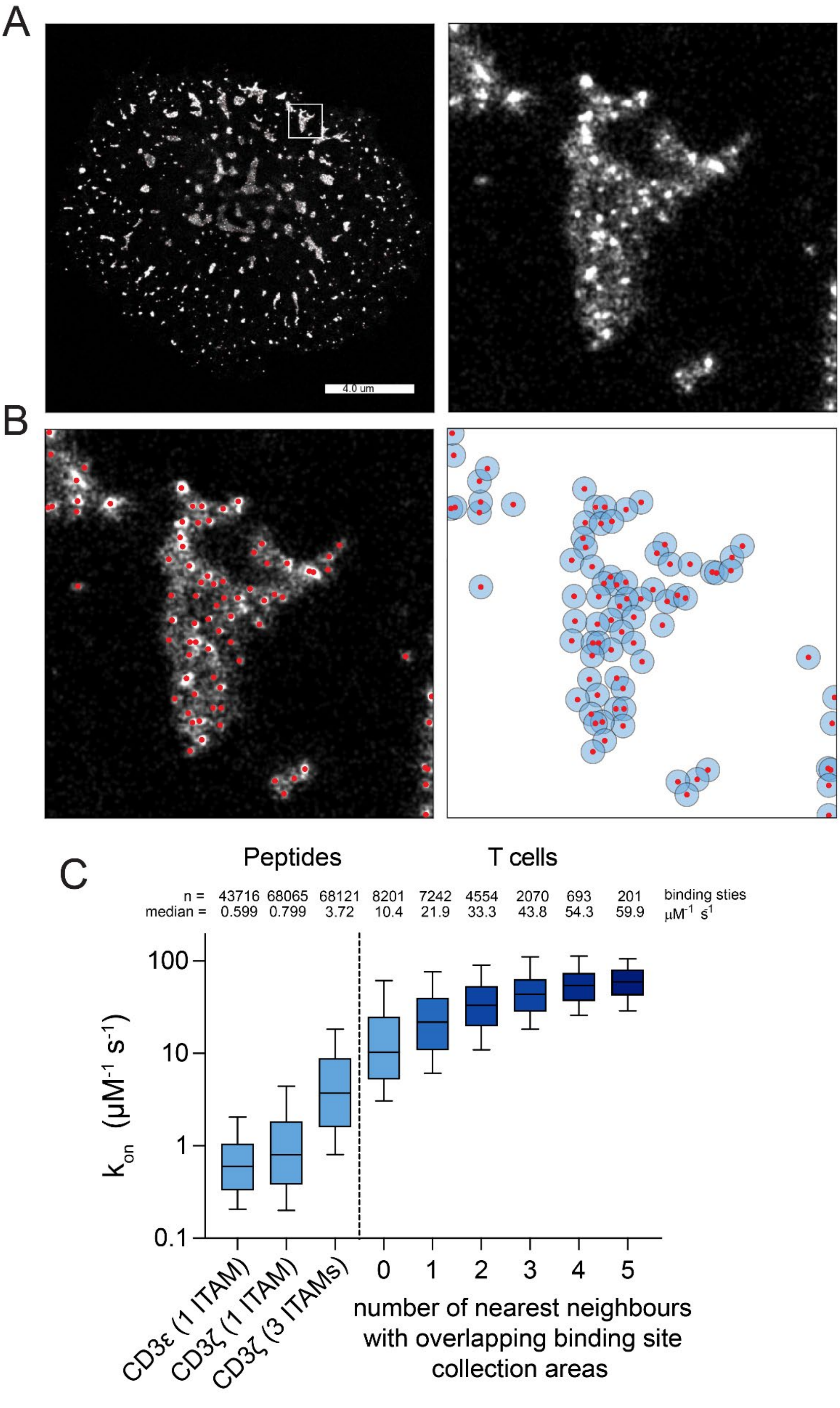


**Figure S4: Estimate of precision and resolution in protein PAINT data.**

(A) Raw reconstructed image of Zap70 binding events in CTLs activated with cognate pMHC. A 1 µm zoomed region is shown on the right.

(B) Zoomed region in (A) overlayed with red dots indicating binding sites identified with the 2D gaussian fitting algorithm on the left, and binding sites with blue circles indicating the radius of binding events collected as belonging to each binding site. Circles are semi-transparent and areas of overlap are evident as darker blue regions.

(C) Binding association rates ($k_{on}$) derived from protein-PAINT measurements for peptides on glass and pTCR binding sites in T cells from the dataset shown in Fig. 1. Peptides immobilized on glass were imaged at low surface density, resulting in well-separated binding sites with minimal overlap between the 45 nm binding-event attribution radii used to group events. In contrast, pTCRs within activated T cells frequently formed clusters, producing overlap between attribution regions of neighboring binding sites. Because binding events within this radius are assigned to each site independently, overlap leads to partial double-counting of events and an increase in the apparent $k_{on}$. Stratifying pTCR binding sites by the number of neighboring sites with overlapping attribution regions revealed a linear scaling of the measured $k_{on}$ with neighbor number, with sites having n neighbors exhibiting approximately n× the association rate of isolated sites. This relationship is consistent with the expected consequences of event sharing between closely spaced sites and supports the ability of the 2D Gaussian fitting approach to reliably identify individual binding sites even in densely packed environments.

Boxes represent interquartile range, with lines indicating median and whiskers representing 10-90 percentile for all identified binding sites in all images. The total number of binding sites from all imaging data and the median $k_{on}$ values are shown above each box plot.

#### Supplementary Figure 5

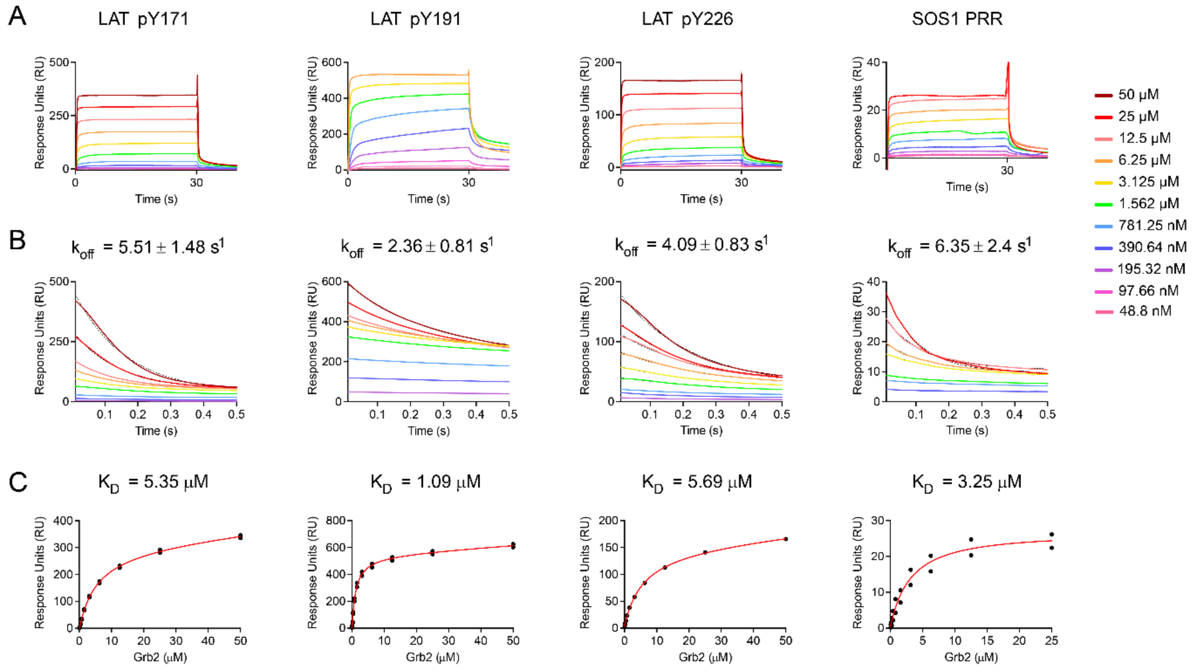


**Figure S5: Binding kinetics of the Grb2-mSG probe measured by SPR.**

(A) Raw overlayed SPR traces of Grb2-mSG probe binding with indicated peptides. Concentrations of probe Zap70 probe injected are shown.

(B) Dissociation constants ($k_{off}$) fit from the dissociation phase of SPR traces for corresponding curves in (A).

(C) Equilibrium binding and $K_D$ fits from data in (A).

**Supplementary Figure 6**

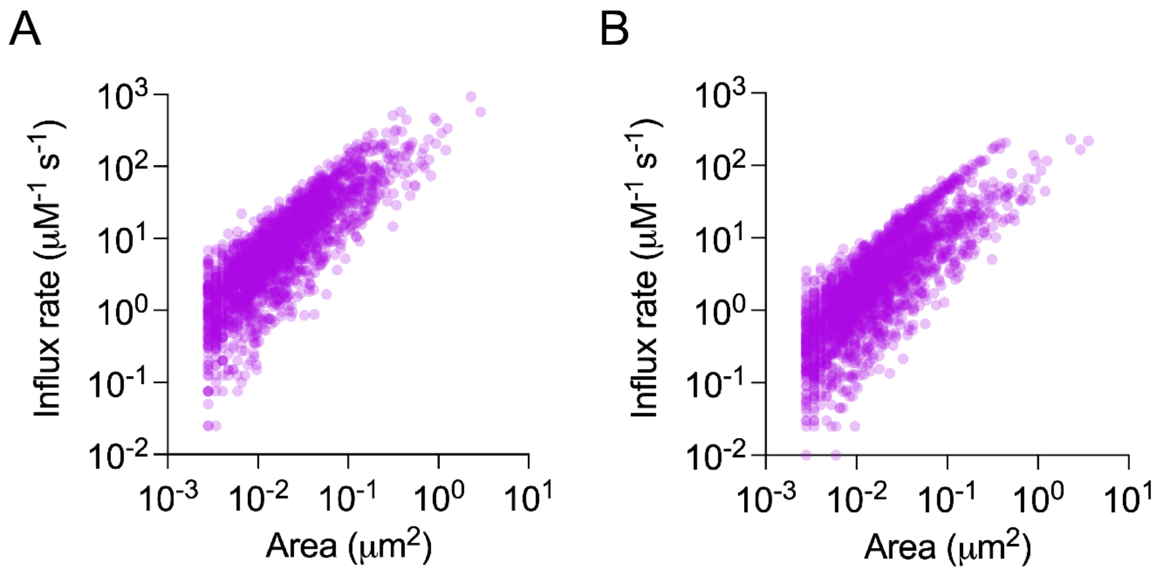


**Figure S6: Binding rates of probes within condensate masked regions.**

(A and B) Influx rates, defined as the rate of new binding event detection for full length Grb2-mSG (A) and Grb2 SH2-mSG (B) protein PAINT probes within a given condensate area (expressed in units of per μM probe per second imaging time) plotted against the area of the condensate. Influx rate is directly proportional to the number of binding sites present[2] within the condensate area, and thus these results indicate a linear relationship between condensate area and the number of pLAT binding sites. Each dot represents one condensate area and data are from the dataset shown in Fig 3 (ie. 448 condensates from 10 cells in 2 biological repeats).

**Supplementary Figure 7**

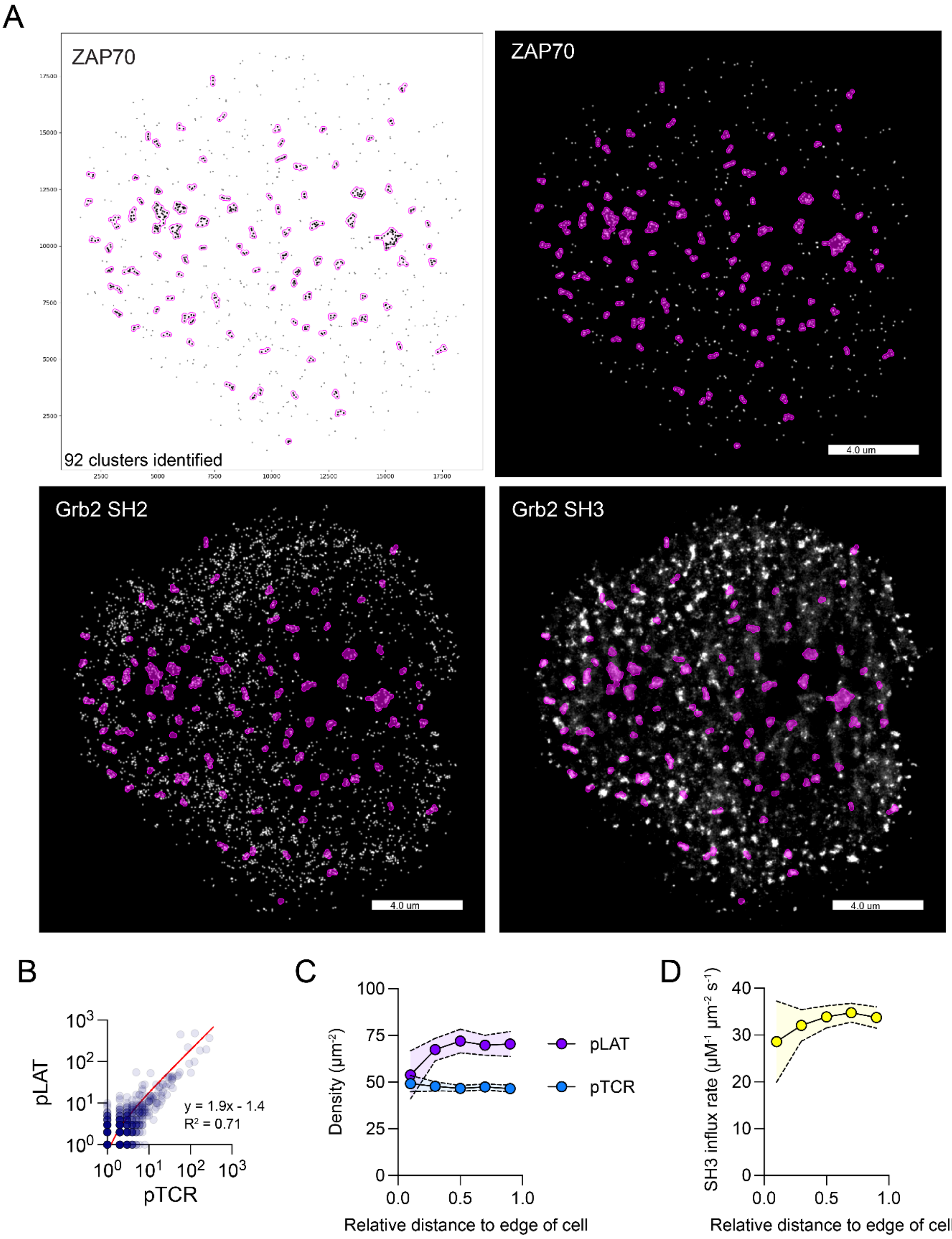


**Figure S7: Defining pTCR clusters.**

(A) Example of identification of clustered pTCR binding regions using DBSCAN. For the DBSCAN algorithm a search radius (ε) of 200 nm and a minimum cluster points of 3 were used. Identified pTCR binding sites (black dots) with DBSCAN-defined clusters (magenta outlines) are shown on the top left. The other subpanels show reconstructed binding event images (binned at 10 nm) with overlayed pTCR binding sites (magenta outlines).

(B) The number of identified pTCR and pLAT binding sites detected within pTCR clustered regions plotted against each other. A linear fit of the data is shown and indicates there are approximately two pLAT molecules for every pTCR within pTCR clustered areas.

(C) Density of pTCR and pLAT within pTCR condensates at different relative distances from the edge of the cell. Density is remarkably consistent for pTCR, and for pLAT, excepting in the bin closest to the edge of the cell in which a subtle but not statistically significant decrease in pLAT is noted.

(D) Influx rate of Grb2 SH3-mNG probe within pTCR cluster regions (expressed in units of per μM probe per area of cluster per second imaging time) plotted against relative distance from the edge of the cell. Influx rates of Grb2 SH3-mNG probe are proportional to the number and density of proline-rich regions in the condensate.

Data are from the dataset shown in Fig 4, with 401 DBSCAN-defined pTCR clusters from 11 cells in 3 biological repeats).
